# Claudin-4 Reconstituted in Unilamellar Vesicles is Sufficient to Form Tight Interfaces that Partition Membrane Proteins

**DOI:** 10.1101/309856

**Authors:** Brian Belardi, Sungmin Son, Michael D. Vahey, Jinzhi Wang, Jianghui Hou, Daniel A. Fletcher

**Affiliations:** Department of Bioengineering and Biophysics Program, University of California, Berkeley, CA, 94720 USA; Division of Biological Systems and Engineering, Lawrence Berkeley National Laboratory, Berkeley, CA, 94720 USA; Department of Internal Medicine & Center for Investigation of Membrane Excitability Disease, Washington University Medical School, St. Louis, MO 63110 USA; Chan Zuckerberg Biohub, San Francisco, CA 94158

## Abstract

Tight junctions have been hypothesized to act as molecular fences in the plasma membrane of epithelial cells, helping to form differentiated apical and basolateral domains. While this fence function is believed to arise from the interaction of four-pass transmembrane claudins, the complexity of tight junctions has made direct evidence of their role as a putative diffusion barrier difficult to obtain. Here we address this challenge by reconstituting claudin-4 into giant unilamellar vesicles using microfluidic jetting. We find that reconstituted claudin-4 is sufficient to form adhesive interfaces between unilamellar vesicles without accessory proteins present in vivo. By controlling the molecular composition of the inner and outer leaflets of jetted membranes, we show that claudin-4-mediated interfaces can drive partitioning of extracellular membrane proteins but not of inner or outer leaflet lipids. Our findings indicate that homotypic interactions of claudins and their small size can contribute to the polarization of epithelial cells.

## Introduction

Nowhere in biology is large-scale membrane organization more apparent than in epithelial tissue, where cell surfaces are differentiated into two distinct domains, the apical and the basolateral (Koichi et al., 1974; Rodriguez-Boulan and Nelson, 1989). The tight junction (TJ), which controls the paracellular flux of ions, solutes, and macromolecules in vertebrate epithelial cells, sits at the boundary between the apical and basolateral membrane domains (Zihni et al., 2016). Due to its unique position, the TJ has long been implicated in the formation of a molecular “fence” between the two domains (Tsukita et al., 2001), which may help to prevent apical-basolateral mixing of proteins and lipids. However, evidence for such a role is conflicting, and whether the TJ acts as a physical barrier to both protein and lipid diffusion in epithelia remains an open question.

Progress in understanding the basic organizational and barrier properties of the TJ has been hindered by its molecular complexity. Work by Van Itallie et al. using proximity ligation and proteomics demonstrated that >400 different proteins reside within the TJ in polarized kidney cells (Van Itallie et al., 2013). This complexity has frustrated efforts to develop a mechanistic picture of putative fence function of the TJ in epithelial cells using traditional top-down cell biological methods, such as knockdowns, knockouts, and knockins. One method to address basic questions about biologically complex systems is to reconstitute key cellular components in vitro and to assay for activity. Such a bottom-up approach has not yet been applied to the TJ.

In a landmark 1998 paper, Tsukita and coworkers uncovered the key membrane protein family necessary for forming intercellular pores at the TJ (Furuse et al., 1998). These proteins, the claudin family, are characterized by a four-pass architecture and are thought to interact across cells through homotypic interactions (Hou et al., 2013). Recently, the first crystal structure of a claudin, claudin-15, revealed one remarkable feature of claudins, namely that the height of their extracellular loops was short, <2 nm (Suzuki et al., 2014). Given the prevailing notion that claudins act as a diffusion barrier at the TJ (Trimble and Grinstein, 2015) and the recent finding that molecular dimensions of adhesive proteins can drive partitioning of non-adhesive biomolecules (Schmid et al., 2016), we sought to test whether claudins alone establish adhesive interfaces between reconstituted membrane vesicles and whether these interfaces are sufficient to organize membranes at physiological length-scales.

Reconstitution of claudin proteins in lipid bilayers provides a direct way to test their interactions and fence function in the absence of other TJ proteins. The central experimental challenge, though, is the lack of methods for inserting oriented multi-pass transmembrane proteins in vesicles large enough for visualizing membrane organization. Here we describe a method for overcoming this challenge using microfluidic jetting. We used this method to study interface formation and membrane partitioning in giant unilamellar vesicles containing a classic claudin, claudin-4 (Cldn4) (Mitic et al., 2003). We found that dense claudin-claudin interfaces form spontaneously between membranes and are sufficient to drive partitioning of extracellular membrane proteins but not of lipids.

## Results and Discussion

Incorporating membrane proteins into giant unilamellar vesicles (GUVs) suitable for fluorescence microscopy is a notoriously difficult problem, with limited in vitro solutions that apply to a small subset of transmembrane proteins (Cole et al., 2015; Dezi et al., 2013; Girard et al., 2004). We decided to tackle the challenge of inserting and orienting Cldn4 in GUVs by making use of microfluidic jetting of black lipid membranes (Richmond et al., 2011; Stachowiak et al., 2008). Cldn4 was selected for incorporation in unilamellar vesicles because it is representative of the largest claudin group, the classic claudins, (Hou et al., 2013) and because it is well-characterized as a barrier claudin in the kidney (Hou et al., 2010; Mitic et al., 2003; Van Itallie et al., 2001). Our strategy for asymmetric incorporation relied on i) fusing proteoliposomes containing purified Cldn4 with lipid monolayers on one side of an infinity chamber (Figure 1A) and ii) biasing the topology of Cldn4 during black lipid membrane formation after the removal of an acrylic divider (Figure 1B). We hypothesized that addition of a large, solubilization tag on one side of the protein would energetically bias the orientation of Cldn4 upon black lipid membrane formation. We, therefore, expressed Cldn4 with an N-terminal GFP tag in yeast cells and purified the protein in DDM detergent (Figure 1-figure supplement 1). Since Cldn4’s N- and C-termini are both located in the cytoplasm, we hypothesized that a large, hydrophilic domain on one side of the protein would restrict the orientation of Cldn4, ultimately placing the GFP tag, and thus the N- and C-termini of the four-pass protein, in the lumen of the vesicle, the equivalent of the cytoplasm in cells.

**Figure 1.**
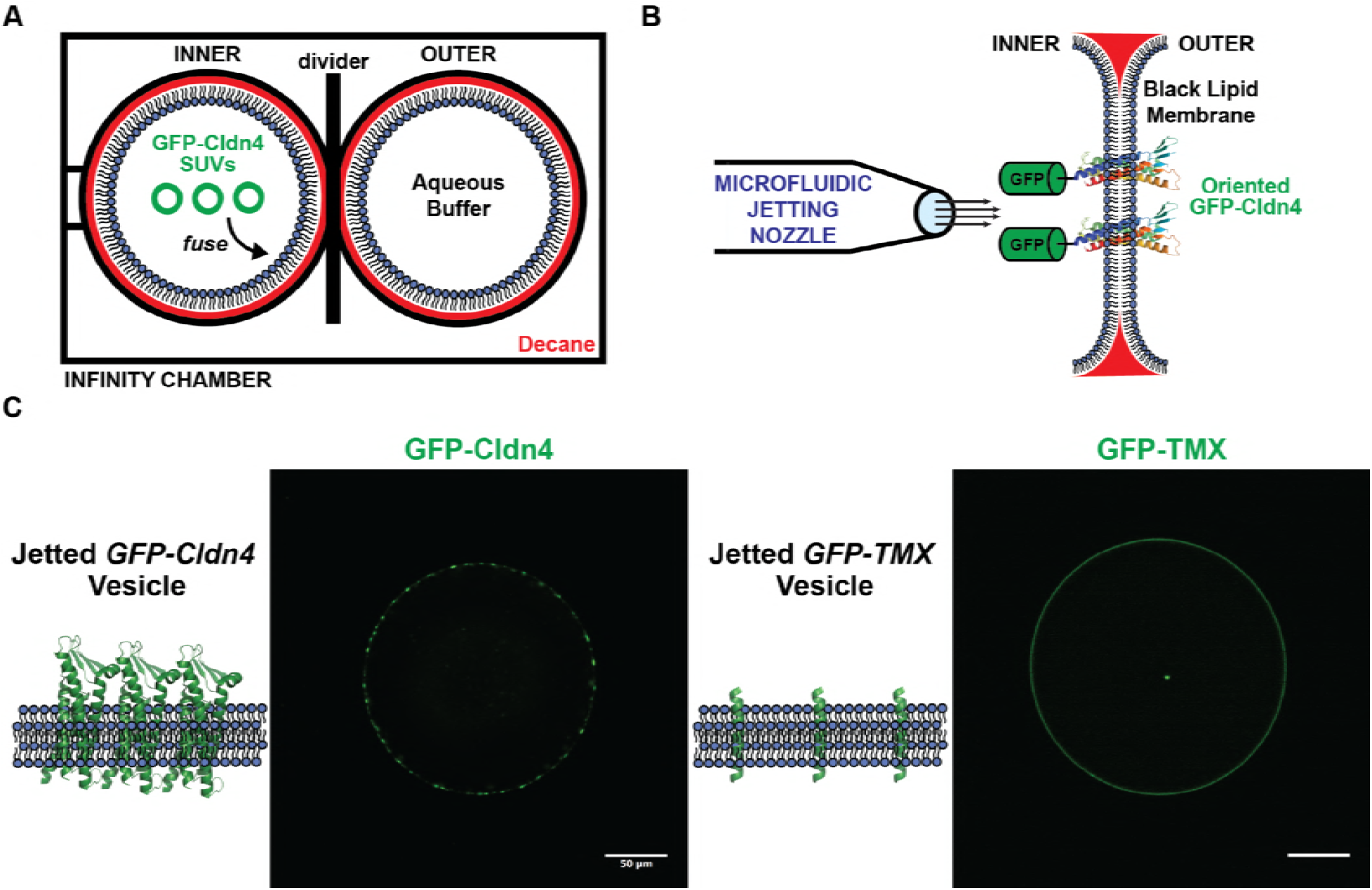
(A) Infinity chamber configuration. To achieve oriented Cldn-4 membrane insertion, GFP-Cldn4 proteoliposomes are incubated in the inner aqueous droplet of an infinity chamber, where they fuse with the DPhPC lipid monolayer at the decane boundary. (B) GFP-Cldn4 black lipid membrane scheme. After the acrylic divider in the infinity chamber is removed, a black lipid membrane spontaneously forms with the hydrophilic portion of Cldn4, i.e. GFP, aligned with the inner aqueous compartment. A microfluidic jetting nozzle is then placed in close proximity to the black lipid bilayer, and GUVs with oriented Cldn4 are jetted by actuating the piezoelectric material. (C) Fluorescent micrographs of single jetted vesicles containing either GFP-Cldn4 or GFP-TMX. Cldn4 organizes into large clusters along the membrane of GUVs.

To generate GUVs with oriented Cldn4, we first formed proteoliposomes with 1,2-diphytanoyl-*sn*-glycero-3-phosphocholine (DPhPC) and purified GFP-Cldn4 using established protocols, i.e. detergent-assisted insertion and Biobead dialysis (Rigaud et al., 1995). We then added GFP-Cldn4 proteoliposomes to either the inner or the outer DPhPC-stabilized droplet of an asymmetric two-droplet infinity chamber. After proteoliposomes had fused with the DPhPC monolayer, the central acrylic divider of the infinity chamber was removed and a black lipid membrane formed spontaneously. We next placed a 25-μm-diameter microfluidic nozzle close to the black lipid membranes and, upon triggering the piezoelectric actuator, formed GUVs by jetting. To test whether GFP-Cldn4 was oriented properly, we incubated both sets of GUVs with Proteinase K, an enzyme capable of digesting exposed GFP (Lorenz et al., 2006). Only in the case where proteoliposomes were added to the outer chamber (corresponding to external orientation of the large, hydrophilic domain) did the GFP signal along the membrane reduce to background levels (Figure 1-figure supplement 2). This experiment suggests that the GFP tag on Cldn4’s N-terminus is unable to traverse the black lipid membrane and can guide Cldn4 into an oriented topology in jetted GUVs.

Next, we imaged GUVs that were generated by incubating GFP-Cldn4 proteoliposomes in the inner droplet of the infinity chamber. This geometry places the extracellular loops of Cldn4 on the outside of the vesicle and the N- and C-termini on the interior of the vesicle. First, we tested if Cldn4 would cluster in the absence of an opposing membrane, as several groups have reported *cis* claudin oligomers in cells (Itallie et al., 2011; Kaufmann et al., 2012; Koval, 2013; Piontek et al., 2011; Rossa et al., 2014). Confocal microscopy showed that GFP-Cldn4 formed large and highly variable clusters in DPhPC GUVs as evidenced by the bright puncta along the jetted membrane (Figure 1C). To test if this clustering was specific to Cldn4, we compared Cldn4’s organization to the membrane distribution of a synthetic transmembrane domain, TMX (Wimley and White, 2000), that should remain monomeric in lipid bilayers. After preparing TMX-GFP proteoliposomes and jetting TMX-GFP GUVs, we observed a uniform distribution of TMX along the lipid membrane (Figure 1C). Our data lends support to the model that claudins interact in a *cis* configuration in lipid bilayers, even in the absence of other TJ adapter proteins.

We then investigated whether claudins in one membrane were able to form adhesive interactions with claudins in a closely apposed membrane. While claudin proteins have long been thought to interact in *trans* across epithelial cells, whether TJ claudins alone are sufficient for interface formation has not yet been examined experimentally. To test this, we incubated Cldn4 proteolipsomes in the inner droplet of an infinity chamber, as described above. After black lipid membranes were formed, we jetted multiple GUVs in quick succession and used confocal microscopy to image the assembly of GUVs. In contrast to the lack of interaction between GUVs in the absence of Cldn4 (Stachowiak et al., 2009), we observed adhesion between GUVs and enrichment of oriented GFP-Cldn4 at GUV-GUV interfaces (Figure 2A). Interfaces appeared to be heterogeneous with respect to Cldn4 distribution; for instance, regions with GFP-Cldn4 puncta and with uniform GFP-Cldn4 co-existed within single interfaces. One striking feature of the GUV assemblies was the absence of close membrane contact at tri-vesicle junctions. In epithelial cells, a specific membrane protein, tricellulin, is necessary for forming tri-cellular TJ contacts (Ikenouchi et al., 2005). It appears from our experiments that Cldn4 is sufficient to form GUV-GUV interfaces but is not capable of forming tri-vesicle contacts, a finding consistent with the localization and function of TJ proteins, including the claudins, in polarized epithelial cells.

**Figure 2.**
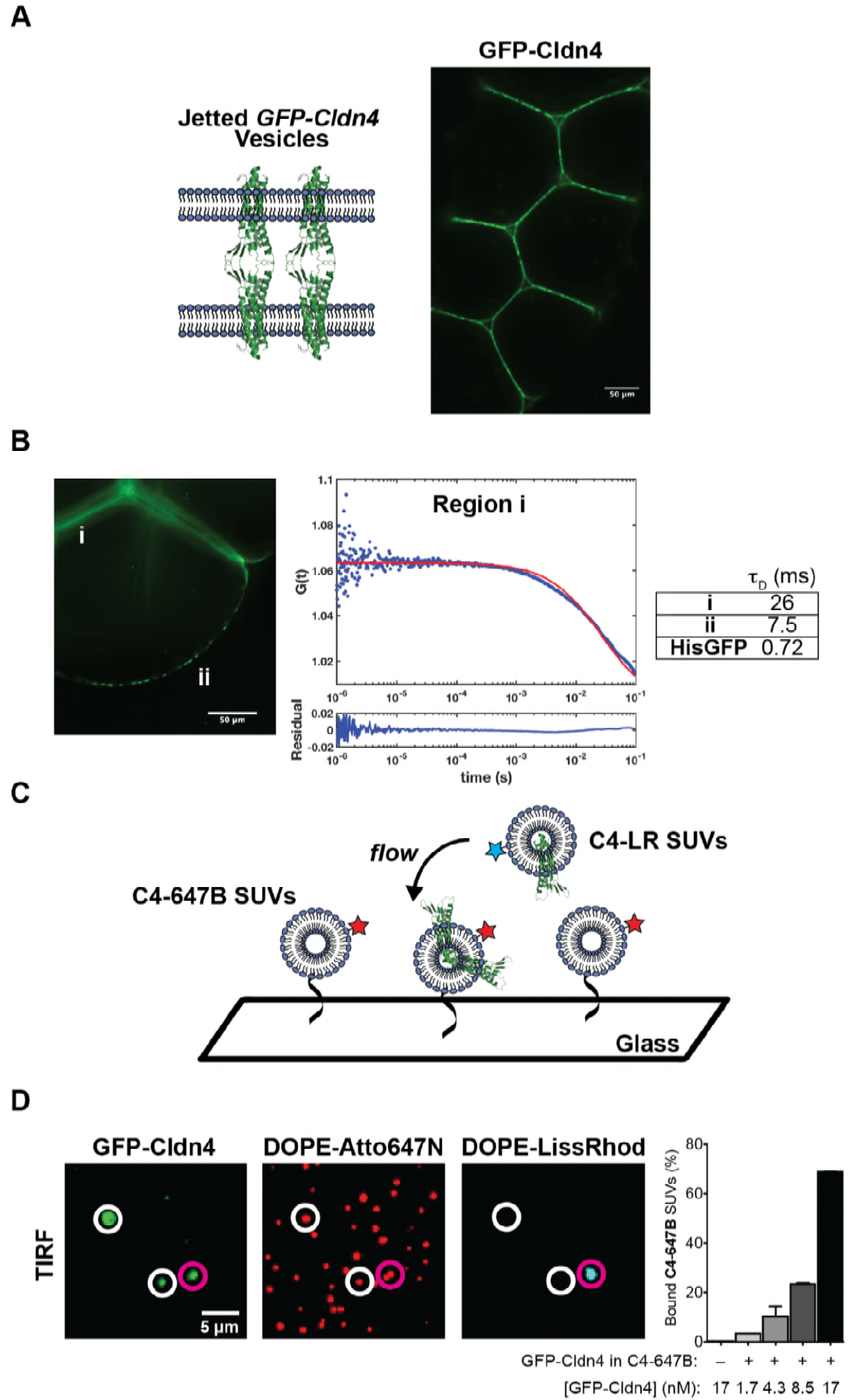
(A) Fluorescent micrographs of jetted GUV assemblies containing GFP-Cldn4. GFP-Cldn4 is enriched at GUV-GUV contacts but not at tri-vesicle junctions. (B) FCS analysis of GFP-Cldn4 in jetted vesicles. Autocorrelation curves were constructed for GFP-Cldn4 at interfaces, i, and at free membrane regions, ii (left). Diffusion times (right, mean, N = 6 for all conditions) were then calculated for the different regions. His-tagged GFP’s diffusion time was also determined in jetted GUVs as a reference. Interfacial GFP-Cldn4 has reduced mobility compared to GFP-Cldn4 at exposed membranes. (C) Single SUV binding assay. C4-647B SUVs are immobilized on a glass slide, and suspended C4-LR SUVs are then applied to immobilized SUVs. Co-localization of the two fluorescent lipids indicates that binding between SUVs due to Cldn4 homotypic adhesion has occurred. (D) TIRF micrographs and quantification of single SUV binding assay. White circles highlight C4-647B SUVs positive for GFP-Cldn4 but negative for C4-LR SUV binding. Magenta circles highlight bound SUVs. Only SUVs positive for GFP-Cldn4 show binding activity (right), demonstrating that reconstituted GFP-Cldn4 can mediated interface formation between SUVs (mean ± SD, N = 13, 9, 13, 12, and 10, with n > 500 particles counted per sample).

We next turned our attention to Cldn4’s dynamics in free membrane regions and in GUV-GUV interfaces. The dynamics of another classic claudin, claudin-1, have previously been characterized in polarized epithelial cells (Shen et al., 2008). In the context of cell-cell contacts, claudin-1’s recovery was found to be slow and accompanied by a large immobile fraction (~78%), suggesting that claudins are statically captured and densely packed at the TJ. We wanted to see if the dynamics of Cldn4 were similar in vitro, even in the absence of other TJ proteins and an underlying cortical cytoskeleton. Fluorescence correlation spectroscopy (FCS) was employed to detect the dynamics of GFP-Cldn4 in two membrane regions of the jetted GUVs, at GUV-GUV interfaces (i) and at free membrane regions (ii) (Figure 2B). We found significantly different autocorrelation curves for the two regions, indicative of different diffusive behavior of claudin-4 in the two regions (Figure 2-figure supplement 1). As our FCS measurements are based neither on a strict 2-D nor a 3-D configuration, and since large protein clusters are known to give rise to anomalous diffusion (Feder et al., 1996), we relied on the model-independent diffusion time parameter to quantify and compare mobility. Higher diffusion times indicate mobility has been significantly reduced. For region A, the diffusion time was 26 ± 4.1 ms (mean ± SD), whereas the diffusion time for region B was 7.5 ± 2.5 ms (Figure 2B), reflecting the fact that interfacial GFP-Cldn4’s mobility had been reduced by a factor of ~3 at membrane interfaces, a result consistent with claudin-1’s dynamics in cells. However, in contrast to in vivo studies, GFP-Cldn4 remained mobile at interfaces, which points to the role other TJ proteins play in forming static claudin structures in epithelial cells. To compare GFP-Cldn4’s diffusion to a known protein, we jetted vesicles containing DOGS-Ni-NTA and complexed His-GFP to the outer leaflet. We found His-GFP’s diffusion time in jetted vesicles to be 0.72 ± 0.56 ms, demonstrating that in free membrane portions of jetted vesicles GFP-Cldn4 with transmembrane domains embedded in the DPhPC lipid bilayer formed clusters with reduced mobility compared to membrane-bound proteins.

Membrane interfaces in cells occur over a range of length scales, and as such, we wanted to test whether Cldn4’s adhesive behavior was dependent on the large surface area presented by GUVs. To quantify Cldn4’s *trans* binding activity at smaller interfaces, we developed a single vesicle binding assay (Figure 2C) based on the GFP-Cldn4 proteoliposomes (100’s of nm in diameter) outlined above. Two different populations of GFP-Cldn4 small unilamellar vesicles (SUVs) were prepared, one containing fluorescent DOPE-Lissamine Rhodamine B (C4-LR SUVs) and the other containing fluorescent DOPE-Atto647N and DOPE-biotin (C4-647B SUVs). Single C4-647B SUVs were tethered to glass surfaces through streptavidin and imaged using total internal reflection microscopy (TIRF). GFP-Cldn4 signal from diffraction-limited, Atto647N-positive particles was only observed for a sub-population of C4-647B SUVs (Figure 2D), which is expected for the detergent-assisted insertion method (Mathiasen et al., 2014). After surface immobilization, we subsequently flowed in C4-LR SUVs and used TIRF to quantify the extent of co-localization (see Supporting Information) of the fluorescent dyes, Lissamine Rhodamine B and Atto647N, as a proxy for Cldn4 binding. First, we noted that only particles positive for GFP-Cldn4 co-localized with C4-LR SUVs (Figure 2D). Second, we found a concentration-dependent increase in bound C4-LR SUVs. No bound C4-LR SUVs were observed in the absence of C4-647B SUVs (Figure 2D). Based on these results, purified GFP-Cldn4 appears competent for *trans* homotypic binding in small as well as large membrane interfaces.

Could claudin’s ability to establish interfaces between membranes, as our data above show for both proteoliposomes and jetted vesicles, contribute to the fence function of tight junctions in the absence of other TJ proteins? Recently, our lab has shown that the molecular dimensions of adhesive proteins can lead to exclusion of non-binding proteins from synthetic interfaces between GUVs (Schmid et al., 2016). We wondered whether this type of exclusion is also relevant to the TJ, since the extracellular portion of claudin was shown to be remarkably short in height by crystallography (Suzuki et al., 2014), and whether in vitro quantification of fence function could help to address continued debate about the TJ’s role in segregating apical and basolateral surfaces. Early work on the TJ pointed to a barrier able to prevent the diffusion of apical components, both proteins and lipids, to the basolateral surface of polarized epithelial cells (Diamond, 1977). Refinements to this model were subsequently made in light of further findings: only outer leaflet lipids appeared to experience a barrier (Dragsten et al., 1981; van Meer and Simons, 1986) and the bulky glycoproteins and glycolipids were especially affected by a fence (Spiegel et al., 1985). After their discovery as the functional transmembrane unit of the TJ (Furuse et al., 1998), claudins were accordingly assigned the role of diffusion barrier between apical and basolateral domains (Trimble and Grinstein, 2015). However, recently, one paper provided evidence of cell polarity in isolated single cells, suggesting that TJs are not needed to partition membrane components (Baas et al., 2004), and in other work, mammary epithelial cells lacking claudin strands appeared to display signs of polarity (Umeda et al., 2006). Due to these contradictory findings, we examined Cldn4-mediated interfaces formed between jetted vesicles for evidence of protein and lipid partitioning.

We jetted GFP-Cldn4 GUVs from two different black lipid membrane configurations. In the first configuration, the inner droplet was incubated with GFP-Cldn4 proteoliposomes containing the far-red fluorescent lipid, DOPE-Atto647N, while the outer droplet was incubated with liposomes containing a red fluorescent lipid, DOPE-LissRhod. In the second configuration, the inner droplet was incubated with GFP-Cldn4 proteoliposomes as before, and the outer droplet was incubated with liposomes containing DOGS-Ni-NTA lipid. For the first configuration, black lipid membranes were formed and again GUVs were jetted in rapid succession, generating vesicles with different fluorophores in the inner and outer leaflets of the GUVs. Following Cldn4-mediated assembly, we did not observe segregation of the two different lipid-fluorophores from interfaces, indicating that Cldn4 alone does not exclude lipids, on either leaflet, from intermembrane junctions (Figure 3A,B). With the second configuration, we also formed black lipid membranes, but before jetting, we incubated the outer leaflet with either of two different sized, His-tagged, nonbinding proteins, one 5 nm in height (5nm-NBP) and the other 15 nm (15nm-NBP), based on different number of mCherry modules (1 and 3, respectively) (Schmid et al., 2016). We then jetted multiple vesicles and imaged the resulting assemblies. For vesicles containing either the 5 or the 15 nm proteins, we observed dramatic segregation of the non-binding protein at Cldn4-mediated interfaces, with complete exclusion for the 15 nm protein (Figure 3A,B). Notably, this is the first in vitro example of an adhesion protein, namely claudin, forming a membrane interface capable of extensive exclusion of a 5 nm protein. These results, therefore, suggest that claudins alone may be able to partition most apical- and basolateral-domain proteins, especially bulky, apical glycoproteins. Our data may also help explain the spatial separation between the TJ and the other epithelial junctions, i.e. adherens junction and desmosomes, which are formed by adhesive proteins with large molecular dimensions, e.g. E-cadherin and desmogleins, respectively.

**Figure 3.**
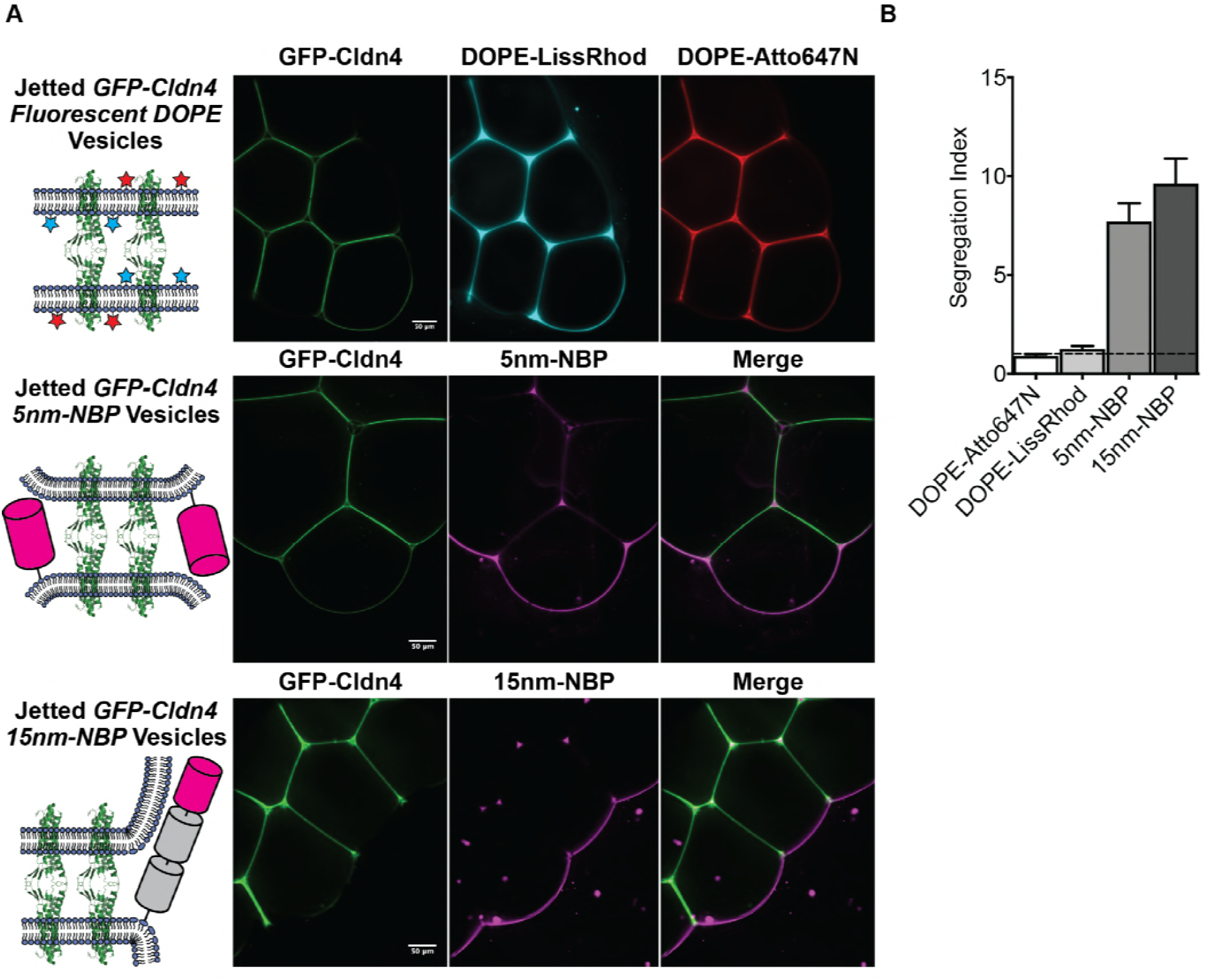
(A) Cldn4-Cldn4 interfaces segregate membrane proteins but not lipids of either leaflet. GFP-Cldn4 GUVs were jetted from two different configurations. In the first configuration, proteoliposomes containing GFP-Cldn4 and DOPE-Atto647N were incubated in the inner droplet, and DOPE-LissRhod liposomes were incubated in the outer droplet. Fluorescent micrographs of GUVs jetted from this configuration (upper panel) do not show evidence of lipid segregation at Cldn4 interfaces. In the second configuration, proteoliposomes containing GFP-Cldn4 were incubated in the inner droplet and liposomes containing DOGS-NiNTA were incubated in the outer droplet. Before jetting, either a His-tagged 5nm-NBP or a 15nm-NBP was applied to the black lipid membrane. Fluorescent micrographs of GUVs jetted from this configuration (middle and lower panels) show that small and large proteins are excluded from Cldn4 interfaces. (B) Quantification of extent of segregation for inner leaflet lipids, DOPE-Atto647N, outer leaflet lipids, DOPE-LissRhod, and non-binding proteins of different molecular lengths, 5nm-NBP and 15nm-NBP. Segregation Index = <Fluorescence Intensity_free membrane_/( Fluorescence Intensity_interface_/2)> (mean ± SD, N = 5 for all conditions).

In summary, we have reconstituted claudin-mediated membrane interfaces in giant vesicles using microfluidic jetting of black lipid membranes. By engineering the molecular architecture of the Cldn4 protein and controlling the geometry of membrane formation prior to jetting, we were able to achieve oriented insertion of a four-pass transmembrane protein into GUVs. With this system, we were able to address one aspect of a longstanding question in epithelial biology, namely whether TJ components form a physical barrier in polarized epithelial tissue. By chemically defining the molecular environment at membrane interfaces in vitro, we have found that Cldn4 can join GUVs together to build assemblies with unique properties. Our data point to a model where the molecular dimensions of claudin proteins limit the ability of proteins on the scale of 5 nm and larger to translocate past the interface, but do not limit lipids embedded in either leaflet. In vivo, this physical segregation model of protein partitioning may be further augmented to restrict lipid mixing through the formation of claudin strands. The highly organized, stable claudin strands assembled by TJ proteins could possibly act as a barrier to lipid diffusion due to claudin membrane density, and thus excluded-volume effects, in epithelial lipid bilayers. The ability to control the orientation of claudin proteins in giant unilamellar vesicles reported here opens up possibilities for more complex reconstitutions and questions, such as examining how TJ plaque proteins, for example the zonula occludens (ZO) proteins, tune the partitioning of membrane lipids and proteins at interfaces.

## Acknowledgements

This work was supported by grants from the NIH (R01GM114344 and R01DK084059). B.B. was supported by the NIH Ruth L. Kirschstein NRSA fellowship from the NIH (1F32GM115091). S.S. was supported by a fellowship from the Life Sciences Research Foundation. M.D.V. was supported by a CASI fellowship from the Burroughs Wellcome Fund. D.A.F. is a Chan Zuckerberg Biohub Investigator.

## Figure Supplements

**Figure 1-figure supplement 1.**
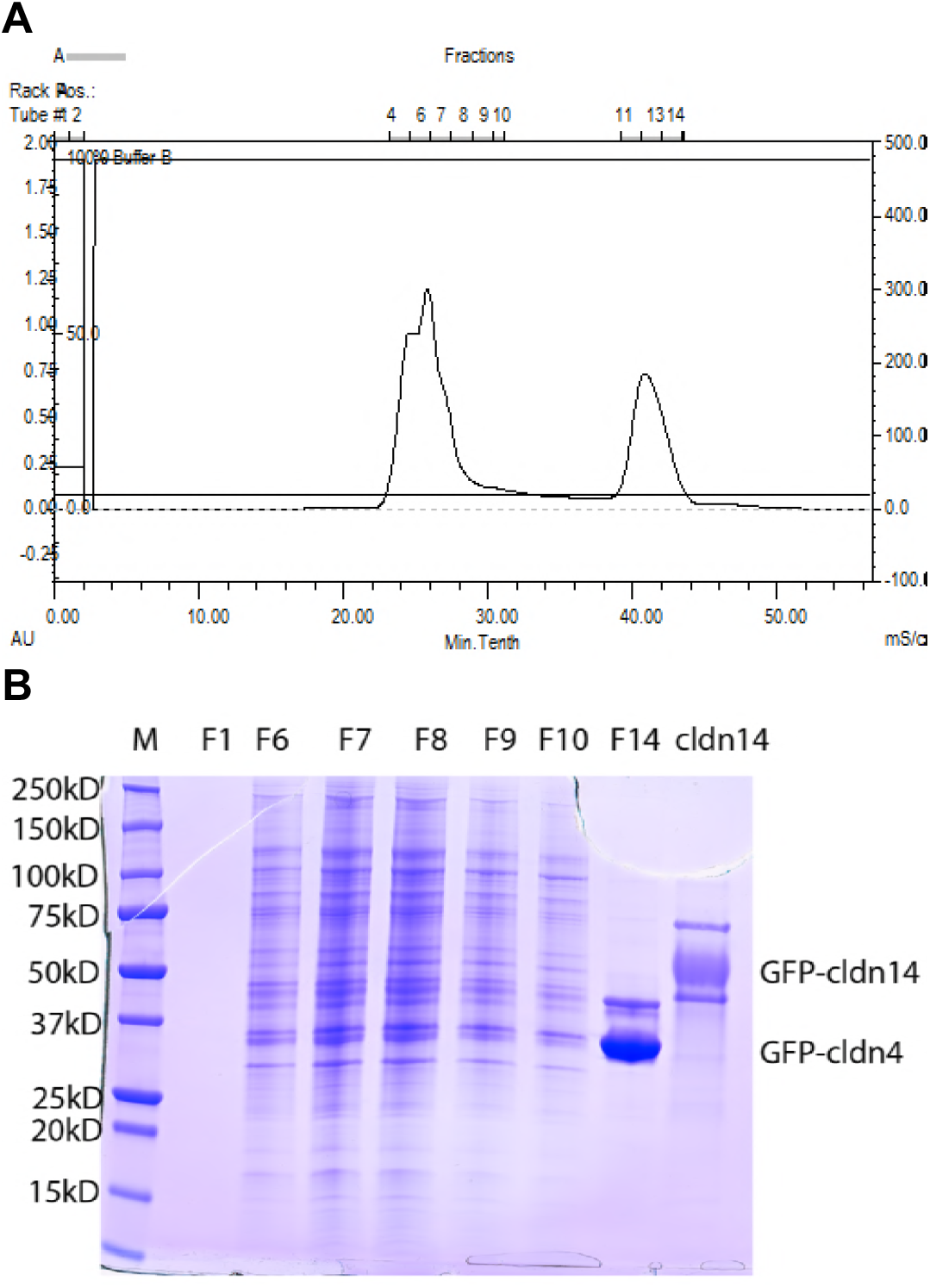
Purification of recombinant GFP-Cldn4 from *Pichia* yeast cells. (**A**) Gel filtration chromatography showing the separation of GFP-Cldn4 proteins (fraction 11-14) from yeast cell endogenous proteins (fraction 4-10). (**B**) SDS-PAGE gel electrophoresis showing the enrichment of GFP-Cldn4 proteins in fraction 14 (F14). A positive control of purified GFP-CLDN14 is also shown (right).

**Figure 1-figure supplement 2.**
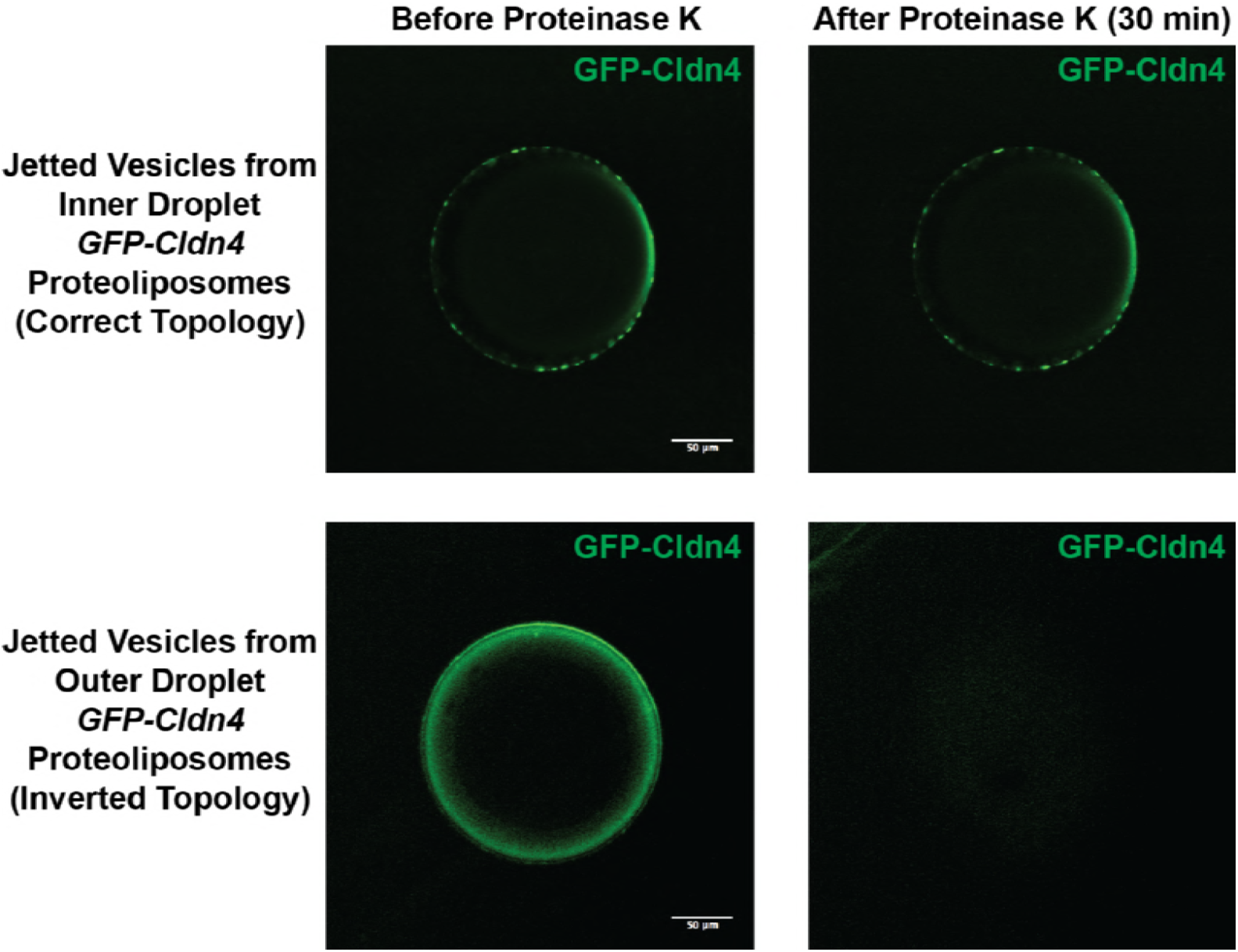
Proteinase K treatment of GFP-Cldn4 GUVs. GFP-Cldn4 GUVs (left) were jetted from black lipid membranes prepared by incubating either the inner droplet with GFP-Cldn4 proteoliposomes (top panel), leading to GFP in the lumen of the vesicle, or the outer droplet with GFP-Cldn4 proteoliposomes (bottom panel), leading to GFP on the exterior of the vesicle. Fluorescence micrographs of the GFP-Cldn4 GUVs show that only the GFP-Cldn4 incubated in the outer droplet is digested by Proteinase K (right), leading to diminished fluorescence.

**Figure 2-figure supplement 1.**
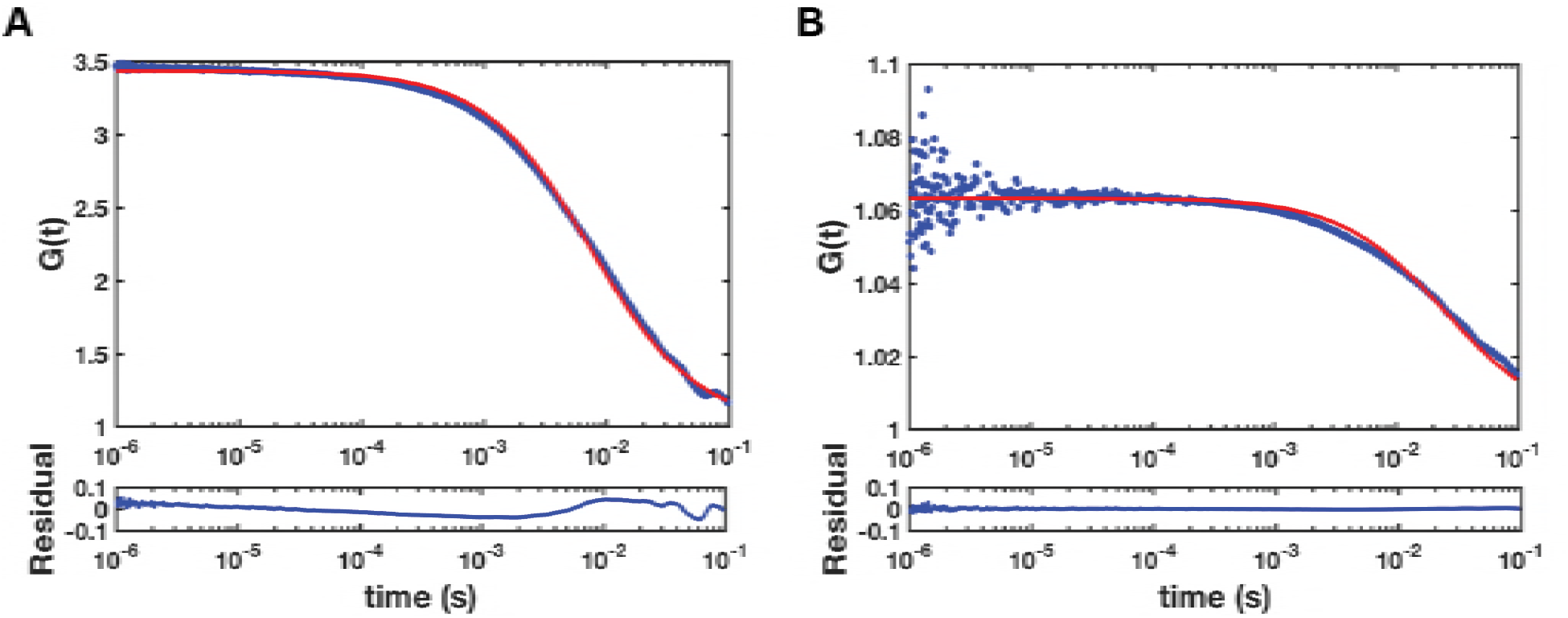
FCS of GFP-Cldn4 in jetted GUVs. (A) Autocorrelation curve and fit for GFP-Cldn4 at free membrane regions. (B) Autocorrelation curve and fit for GFP-Cldn4 at membrane interfaces. Repeated from Figure 2B for comparative purposes. The diffusion time for GFP-Cldn4 is reduced by over a factor of 3 at membrane interfaces compared to free membrane regions.

## Materials and Methods

### General methods

All of the chemical reagents were of analytical grade, obtained from commercial suppliers, and used without further purification, unless otherwise noted. Proteinase K was purchased from New England Biolabs. 1,2-diphytanoyl-*sn*-glycero-3-phophoscholine (DPhPC), 1,2-dioleoyl-*sn*-glycero-3-phosphocholine (DOPC), 1,2-dioleoyl-*sn*-glycero-3-[(N-(5-amino-1-carboxypentyl)iminodiacetic acid)succinyl], nickel salt (DOGS-Ni-NTA), 1,2-dioleoyl-*sn*-glycero-3-phosphoethanolamine-N-(cap biotinyl), sodium salt (DOPE-biotin), 1,2-distearoyl-*sn*-glycero-3-phosphoethanolamine-N-[biotinyl(polyethyleneglycol)-2000], ammonium salt (DSPE-PEG-biotin), 1,2-dioleoyl-*sn*-glycero-3-phosphoethanolamine-N-[methoxy(polyethylene glycol)-2000], ammonium salt (DOPE-PEG) and 1,2-dioleoyl-sn-glycero-3-phosphoethanolamine-N-(lissamine rhodamine B sulfonyl), ammonium salt (DOPE-Lissamine Rhodamine B) were obtained from Avanti Polar Lipids, Inc. 1,2-Dioleoyl-sn-glycero-3-phosphoethanolamine labeled with Atto 647N (DOPE-Atto647N) was purchased from Sigma Aldrich.

Fluorescence imaging was carried out on a Ti Eclipse microscope (Nikon) equipped with a CSU-X spinning disk confocal module (Yokogawa) and a Zyla sCMOS camera (Andor). Fluorescence micrographs of giant vesicles were acquired with either a 10x (Nikon, NA 0.3) or a 20x objective (Nikon, NA 0.45). TIRF imaging was performed on the Ti Eclipse microscope (Nikon) using a 60x objective (Nikon, NA 1.49 TIRF) and an iXon Ultra Em-CCD camera (Andor).

Fluorescence correlation spectroscopy (FCS) data sets were acquired on a custom-built set-up with an inverted Nikon Eclipse TE2000 microscope (Nikon) equipped with a 100x oil immersion objective (Nikon, NA 1.49 TIRF) and a 488 nm CW laser as an excitation source (Sapphire, Coherent). Several turning mirrors were used to center and shape the laser as it was focused into a solution of Atto488. Fluorescence emission was collected on an avalanche photodiode (Excelitas, SPCM-AQR-14), and photon arrival times were measured with the counter module of an NI DAQ board (National Instruments, PCI-6321). Six 20 s photon streams were collected for each position on GUVs. Photon data were analyzed using Matlab scripts (Mathworks).

### Protein Expression and Purification

The cDNA encoding full-length human claudin-4 (Cldn-4) was cloned into the pBV-3 vector in frame with the gene encoding the GFP protein to create a N-terminal fusion protein. The construct was expressed in *Pichia pastoris*. Cells were disrupted by milling (Retsch MM400) and resuspended in lysis buffer containing 50 mM Tris, pH 8.0, and 500 mM NaCl. Lysate was extracted with 2% (w/v) *n*-dodecyl β-d-maltopyranoside (DDM, Anatrace) for 2 h with stirring at 4 °C and then centrifuged for 1 h at 30,000xg. Supernatant was added to cobalt-charged resin (G-Biosciences), and the suspension was mixed by inversion for 3 h. Resin was then washed with ten column volumes of buffer containing 20 mM Tris, pH 8.0, 500 mM NaCl, 10 mM imidazole, 4 mM DDM and eluted with buffer containing 250 mM imidazole. The eluted proteins were concentrated and further purified by gel filtration using a Superose 6 column (GE Healthcare) pre-equilibrated with buffer containing 20 mM Tris, pH 8.0, 150 mM NaCl, 5 mM dithiothreitol (DTT), and 1 mM DDM. Peak fractions corresponding to the monomeric claudin-4 proteins were collected and concentrated to ~8.5 μM.

TMX, a synthetic transmembrane domain bearing the sequence ‘WNALAAVAAALAAVAAALAAVAA,’ was selected for its reported ability to insert into lipid membranes (Wimley and White, 2000). We cloned TMX into the bacterial expression vector pRSETa with a C-terminal eGFP fusion and an N-terminal fusion consisting of a 6x His tag, a maltose binding protein (MBP), and a TEV cleavage site. Both N-terminal MBP and C-terminal eGFP fusions enhance the solubility of the intervening hydrophobic stretch, allowing us to express it in the cytoplasm of E. coli. DNA fragments for each of these modules were amplified using PCR (MBP, eGFP) or synthesized as a gBlock fragment (TMX) and inserted into pRSETa between NheI and EcoRI sites. For protein production, 1 – 2L of cells were grown to an OD_600_ of 0.7, induced with 250 μM IPTG and cultured overnight at 16 °C. After 16 h, cells were harvested by centrifugation and resuspended in lysis buffer consisting of 25mM HEPES, pH 7.5, 50 mM NaCl, 0.5 mM TCEP, 1% Triton X-100, and supplemented with PMSF and DNase I. After briefly pulse-sonicating and incubating for 30 minutes at 4 °C, lysed cells were pelleted to remove cell debris and the lysate was circulated over a 5 mL MBPTrap HP column for 2 h. Following binding, the detergent concentration was gradually reduced to 0.1% and bound protein was eluted using 10 mM maltose. Finally, cleavage with TEV protease was performed overnight at 4 °C using a 1:20 molar ratio of TEV to TMX. Following cleavage, residual MBP, TEV, and uncleaved protein is removed by passing the protease-treated mixture over a HisTrap column.

### Infinity Chamber Fabrication

Infinity chambers were fabricated by first cutting 4.5 mm-thick acrylic sheets (McMaster-Carr) using a laser cutter (Versa Laser). The inner and outer chambers were fabricated to be 4 mm in diameter and separated by a 0.15 mm-wide slot. 1.7 mm holes were then drilled into the side of the inner chamber. Subsequently, a thin, 0.2 mm acrylic coverslip was cemented (TAP Plastics, Acrylic Cement) to the inner chamber, and a no. 1.5 glass coverslip coated in poly-L-lysine was glued (Norland Optical Adhesive 60) to the bottom of the outer chamber for imaging. After placing a thin, 0.2 mm acrylic divider between the two chambers, latex (McMaster-Carr) was glued (Gorilla Super Glue) over the inner chamber hole and pierced with a 23G needle.

### Proteoliposome Preparation

Proteolipsomes were generated according to established procedures (Martens et al., 2007). Briefly, GFP-Cldn4 in 20 mM Tris, pH 8.0, 150 mM NaCl, 5 mM DTT, and 1mM DDM was diluted 1:1 with buffer containing 2% Octyl β-D-glucopyranoside (OGP), 25 mM HEPES, pH 7.5, and 150 mM NaCl to a final volume of 80 μL. 20 μL of 10 mM DPhPC SUVs in 25 mM HEPES, pH 7.5, 150 mM NaCl was added to the protein-detergent solution. The mixture was incubated for 15 min at rt. OGP-solubilized DPhPC/GFP-Cldn4 was then diluted four-fold with 25 mM HEPES, pH 7.5, 150 mM NaCl, and 0.5 mM TCEP and dialyzed overnight against buffer containing 25 mM HEPES, pH 7.5, 150 mM NaCl, 0.5 mM TCEP, and 10 g of BioBeads (Thermo). A similar procedure was used to prepare TMX-GFP proteoliposomes and DOPC-based proteoliposomes for the single SUV binding assay.

### Black Lipid Membrane Formation

A planar bilayer between the inner and outer compartments of the infinity chamber was formed by following the protocol developed by Richmond et al (Richmond et al., 2011). In brief, a 15 μL volume of a freshly prepared 25 mg/mL solution of DPhPC in n-decane was first applied to the central acrylic divider in the infinity chamber. Subsequently, 40 μL of a 5-fold dilution of the GFP-Cldn4 proteoliposome solution was added to the inner chamber, and a 40 μL volume of buffer containing 25 mM HEPES, pH 7.5, 150 mM NaCl, and 0.5 mM of TCEP was added to the outer chamber. Solutions in the infinity chamber were incubated for 15 min at rt, 45 min at 4 °C, and finally for 5 min at rt. The acrylic divider was subsequently removed, and after 15 min, oil evacuation and black lipid membrane formation was monitored using brightfield microscopy.

### Microfluidic Jetting of Black Lipid Membranes

GUVs were formed by placing a microfluidic jetting nozzle (Microfab Technologies, Inc., single jet microdispensing device with 25 μm orifice) filled with a 10% OptiPrep (Sigma-Aldrich), 25 mM HEPES, pH 7.5, 150 mM NaCl solution in close proximity, <200 μm, to GFP-Cldn4 planar bilayers. This solution ultimately constitutes the lumen of the jetted GUVs and has a matched osmolarity, but different density, compared to the outer buffer. The piezoelectric actuator was controlled by a waveform generator (Agilent) and an amplifier (Krohn-Hite) with pulse train envelops designed by a custom Matlab script. For jetting of GFP-Cldn4 GUVs, the actuator was triggered by an increasing parabolic envelop defined by 40 trapezoidal bursts at 15 kHz, 3 μs rise and fall times, a 30 μs hold time, and a maximum voltage of 15-30 V. Planar bilayer deformation and GUV formation were monitored using brightfield microscopy with a high-speed camera (Photron, 1024PCI). GUVs formed by microfluidic jetting sunk to the poly-L-lysine-coated coverglass due to the density mismatch between the interior of the GUVs and the surrounding buffer. To determine the orientation of GFP-Cldn4 in GUVs, a volume of 1 μL of 800 units/mL solution of Proteinase K was added to the outer buffer after GUV formation and incubated with jetted GUVs for 30 min.

### Single SUV Binding Assay

Proteoliposomes containing GFP-Cldn4 were prepared as described above. Two separate populations of GFP-Cldn4 proteoliposomes were generated for the single SUV assay: one with a lipid composition of DOPC (99.8%), DOPE-Atto647N (0.1%), and DOPE-biotin (0.1%), and the other with a lipid composition of DOPC (99.9%) and DOPE-Lissamine Rhodamine B (0.1%). Both sets of proteoliposomes were purified and diluted by gel filtration with a S400 HR gel filtration column (GE Healthcare) to remove lipid aggregates before use. To construct a sandwich channel, glass slides were piranha cleaned and treated with poly-L-lysine-g-PEG to passivate the surface. Glass coverslips, on the other hand, were RCA cleaned. The two glass surfaces were adhered to one another with double-sided tape, and an immobile supported lipid bilayer (SLB) was formed on the glass coverslip by incubating the chamber with a 1 mg/mL solution of SUVs containing DOPC (97%), DOPE-PEG (2.94%), and DOPE-PEG-biotin (0.06%) for 10 min. DOPE-PEG acts to passivate the coverslip surface, while DOPE-PEG-biotin acts as tethering points for single SUVs. The channel was washed with a volume of 200 μL of buffer containing 25 mM HEPES, pH 7.5, 150 mM NaCl, and then incubated with a 50 μg/mL solution of streptavidin (Life Technologies) for 5 min. The channel was washed with buffer and then incubated with purified proteoliposomes containing GFP-Cldn4, DOPC, DOPE-Atto647N, and DOPE-biotin for 10 min. The channel was subsequently washed with buffer and then either imaged by TIRF microscopy or incubated with the second set of proteoliposomes containing GFP-Cldn4, DOPC, and DOPE-Lissamine Rhodamine B at various dilutions and followed by washing. To determine the percentage of bound DOPE-Atto647N-containing proteoliposomes, a Matlab script was used. Thresholding was first applied to each fluorescence channel. Fluorescent particles were then located and their centroids were determined using a peak-finding algorithm. Positions of particles positive for both DOPE-Atto647N and GFP-Cldn4 were first saved and then compared to positions of DOPE-Lissamine Rhodamine B-positive particles to calculate the percentage of bound DOPE-Atto647N-containing proteoliposomes. At least ~500 particles/sample were analyzed for each data point.

### Jetted GUV Segregation Assay

For the jetted GUV segregation assay, four separate proteoliposome/liposomes were formulated as described above: proteoliposomes containing GFP-Cldn4/DPhPC, proteoliposomes containing GFP-Cldn4/DPhPC/DOPE-Atto647N (0.1%), liposomes containing DPhPC/DOGS-Ni-NTA (2.5%), and liposomes containing DPhPC/DOPE-Lissamine Rhodamine B (0.1%). Using the fabricated infinity chamber, a 15 μL volume of a freshly prepared 25 mg/mL DPhPC solution in n-decane was applied to the central acrylic divider. A 40 μL volume of a five-fold dilution of proteoliposomes containing either GFP-Cldn4/DPhPC/DOPE-Atto647N or GFP-Cldn4/DPhPC was incubated in the inner chamber, while a 40 μL volume of a five-fold dilution of liposomes containing either DPhPC/DOPE-Lissamine Rhodamine B or DPhPC/DOGS-Ni-NTA was incubated in the outer chamber simultaneously. After the acrylic divider was removed, a black lipid membrane was formed over the course of 15 min. For the fluorescent lipid experiment, GFP-Cldn4 GUVs were jetted in rapid succession with DOPE-Atto647N embedded in the inner leaflet and DOPE-Lissamine Rhodamine B embedded in the outer leaflet. For the fluorescent protein experiment, after planar bilayer formation, a volume of 1 μL of a 20 μM solution of either the 5 nm or the 15 nm His-tagged mCherry protein was added to the outer chamber and incubated for 30 min. GUVs were then generated in rapid succession to produce assemblies of GFP-Cldn4 interfaces.

